# Loss of PTEN promotes formation of signaling-capable clathrin-coated pits

**DOI:** 10.1101/137760

**Authors:** Luciana K Rosselli-Murai, Joel A Yates, Sei Yoshida, Julia Bourg, Kenneth K.Y. Ho, Megan White, Julia Prisby, Xinyu Tan, Megan Altemus, Liwei Bao, Zhi-Fen Wu, Sarah Veatch, Joel Swanson, Sofia Merajver, Allen P Liu

**Author notes:** Correspondence and request for material should be addressed to S.D.M. and A.P.L.

## Abstract

Defective endocytosis and vesicular trafficking of signaling receptors has recently emerged as a multifaceted hallmark of malignant cells. Clathrin-coated pits (CCPs), the fundamental unit of clathrin-mediated endocytosis, display highly heterogeneous dynamics on the plasma membrane where they can take from 20 seconds to over a minute to form cytosolic coated-vesicles. Despite the large number of cargo molecules that traffic through CCPs, it is not well understood whether signaling receptors activated in cancer, such as epidermal growth factor receptor (EGFR), are regulated through a specific subset of CCPs. The signaling lipid phosphatidylinositol (3,4,5)-triphosphate (PI(3,4,5)P_3_), which is dephosphorylated by phosphatase tensin homolog (PTEN), is a potent tumorigenic signaling lipid that is present in excess in many types of cancers. Using total internal reflection fluorescence microscopy and automated tracking and detection of CCPs, we find PTEN and EGF bound EGFR are enriched in a distinct subset of short-lived CCPs that corresponded with clathrin-dependent EGF-induced signaling. By deleting PTEN using CRISPR-Cas9 and reconstituting PTEN, we demonstrate that PTEN plays a role in the regulation of CCP dynamics; this appears to recapitulate CCP dynamics in highly metastatic PTEN-deleted cancer cells where we find a larger proportion of short-lived CCPs and higher initiation density compared to the normal cells. Furthermore, increased PI(3,4,5)P_3_ results in higher proportion of short-lived CCPs, an effect that recapitulates PTEN deletion. Our findings provide strong evidence for the existence of short-lived ‘signaling-capable’ CCPs. Altogether, these findings demonstrate the importance of PTEN and PI(3,4,5)P_3_ in regulating CCP dynamics and assign a new function to PTEN as a modulator of signaling-capable CCPs.

## Introduction

Clathrin-mediated endocytosis (CME) is one of the most important endocytic routes for ligand and receptor uptake. The canonical multi-step process of CME includes its initiation (also known as nucleation), maturation, invagination, and scission of clathrin-coated pits (CCPs) to form single cytosolic vesicles ^1^. In recent years, live cell fluorescence microscopy has enabled researchers to observe the dynamics of CCP assembly in real time. Numerous studies have revealed recruitment of cargo binding adaptor proteins and endocytic accessory proteins during CCP formation ^2–4^.

The dynamics of CCPs is highly heterogeneous, and they can be regulated by cargo, ^5^, cargo-adaptor interactions ^6^, clustering of receptors ^7^, and cytoskeleton/membrane tension ^8–10^. Besides the important roles of protein-protein interactions ^11^, phosphoinositides (PI) are also critical regulators of both intracellular membrane trafficking and cell signaling ^12^. PI levels in a cell are dynamically and spatially regulated by kinases and phosphatases, thereby establishing a code of membrane identity. Among the PIs, PI(4,5)P_2_, PI(3,4)P_2_, and PI(3,4,5)P_3_ are found primarily on the plasma membrane. Notably, PI(4,5)P_2_ is a master regulator of plasma membrane function, including CCP dynamics regulation and membrane-cytoskeleton interactions ^13^. Consequently, lipid modifying enzymes that are not part of the structural CCP core can also influence CME through modulating PI(4,5)P_2_ levels ^14–16^. While the relatively abundant PI(4,5)P_2_ is essential for endocytic vesicle formation ^17–19^, and recent work has also revealed the regulation of CME by PI(3,4)P_2_ ^20^, the role of PI(3,4,5)P_3_ in regulating trafficking is less understood.

Previous work has revealed the existence of a large population of short-lived CCPs that form and abort at the plasma membrane in under a lifetime interval of 20 seconds ^3,21^. A key concept that resulted from this work is that a failure to recruit cargo results in short-lived abortive CCPs. In this manner, cargo recruitment is an input to the endocytic checkpoint for monitoring the fidelity of CCP formation ^21^. Although these short-lived CCPs are the most abundant CCPs at the plasma membrane, whether they have any functional specialization, similar to cargo-dependent control of CCP dynamics for GPCR and transferrin receptor (TfnR) ^5,7^, remains unclear. The low clathrin assembly in short-lived CCPs should have smaller cargo capacity and thus have a higher probability of being abortive. Although some have suggested that a subpopulation of clathrin structures can potentially promote sustained receptor signaling at the plasma membrane ^22,5,23^, the functional significance of short-lived CCPs has never been reported.

As a dynamic plasma membrane remodeling process, CME is highly integrated into signal transduction pathways ^24^. Phosphatase and tensin homolog (PTEN) is a tumor suppressor frequently mutated or inactivated in human cancers, including up to 30–40% of breast cancers, resulting in hyper-activation of the phosphoinositide 3-kinase (PI3K)/Akt signaling pathway ^25,26^. To inhibit cancer-promoting signal transduction, PTEN efficiently antagonizes the PI3K/Akt/mammalian target of rapamycin (mTOR) pathway through its lipid phosphatase activity in response to the activation of receptor tyrosine kinases. PTEN exhibits dual protein and lipid phosphatase activity. The lipid phosphatase activity of PTEN, which is a critical component of its tumor suppressor activity, hydrolyzes PI(3,4,5)P_3_ to PI(4,5)P_2_ and loss of PTEN leads to accumulation of PI(3,4,5)P_3_ at the plasma membrane ^27^. Although PTEN is mainly cytosolic, several studies have revealed transient PTEN recruitment to the plasma membrane to suppress PI(3,4,5)P_3_ signaling ^28–30^, and an engineered PTEN with enhanced plasma membrane activity has been generated ^31^. Given the fact that PI(4,5)P_2_ is known to enhance PTEN association with the plasma membrane to activate PTEN’s phosphatase activity ^32^ and the importance of PI(4,5)P_2_ in CCP formation, it seems reasonable to hypothesize that PTEN may regulate CCP dynamics. Despite the importance of PTEN activity at the plasma membrane, little is known about PTEN function on other cellular membranes. A recent study has shown that PTEN is activated by tethering to early endosomes via association with PI(3)P to inactivate PI3K/Akt signaling ^33^. While PTEN activity and localization have received intense attention in the past few years ^32,34^, how PTEN mechanistically relates to endocytic activity remains poorly understood.

In the present study, we find that clathrin scaffolds are important for EGF-induced signaling at the plasma membrane. Importantly, we find that both EGFR and PTEN are localized to short-lived CCPs. We examined the role of PTEN more closely by reconstituting PTEN in triple negative inflammatory breast cancer SUM149 cells, which do not express functional PTEN due to a genetic micro-amplification within the PTEN gene ^35,36^, and by deleting PTEN in MCF10A and MDA231 cells. By analyzing CCP dynamics, we demonstrate a role for PTEN in regulating the proportion of short-lived CCPs and initiation density. The effect of PTEN is exerted through its lipid phosphatase activity, as directly increasing the concentration of PI(3,4,5)P_3_ in the plasma membrane results in an increase in the number of short-lived CCPs. These findings provide a mechanistic framework for the role of PTEN in CME and reveal a novel function for PTEN in controlling CCP dynamics. Together, our results support the existence of EGF-induced short-lived ‘signaling-capable’ CCPs, which may provide another avenue to target aggressive breast and other PTEN-null cancers.

## Results

### Clathrin inhibition reduces EGF-stimulated Akt phosphorylation in wild type cells but not in PTEN-null cells

EGF binding to EGFR promotes its incorporation to clathrin structures in the plasma membrane concomitantly with initiation of the PI3K signaling cascade ^37^. Since SUM149 cells have endogenously absent PTEN protein expression, we wondered whether formation of CCPs would regulate EGFR signaling in a PTEN-dependent manner. To this end, we first generated PTEN knockout (PTEN-null) MCF10A and MDA231 cell lines using CRISPR-Cas9 by directing Cas9 cleavage to a target site at amino acid position 164 inside the PTEN phosphatase domain (Fig. 1a) and used non-homologous end joining to create mutations resulting in a nonsense mutation and loss of protein expression (Fig. 1b and Sup. Fig. 1a). As expected, deletion of PTEN in both MCF10A and MDA231 cells, led to enhanced cell migration compared to their isogenic counterparts in a wound healing assay (Fig. 1c, d, Sup. Fig. 1b, c).

**Figure 1.**
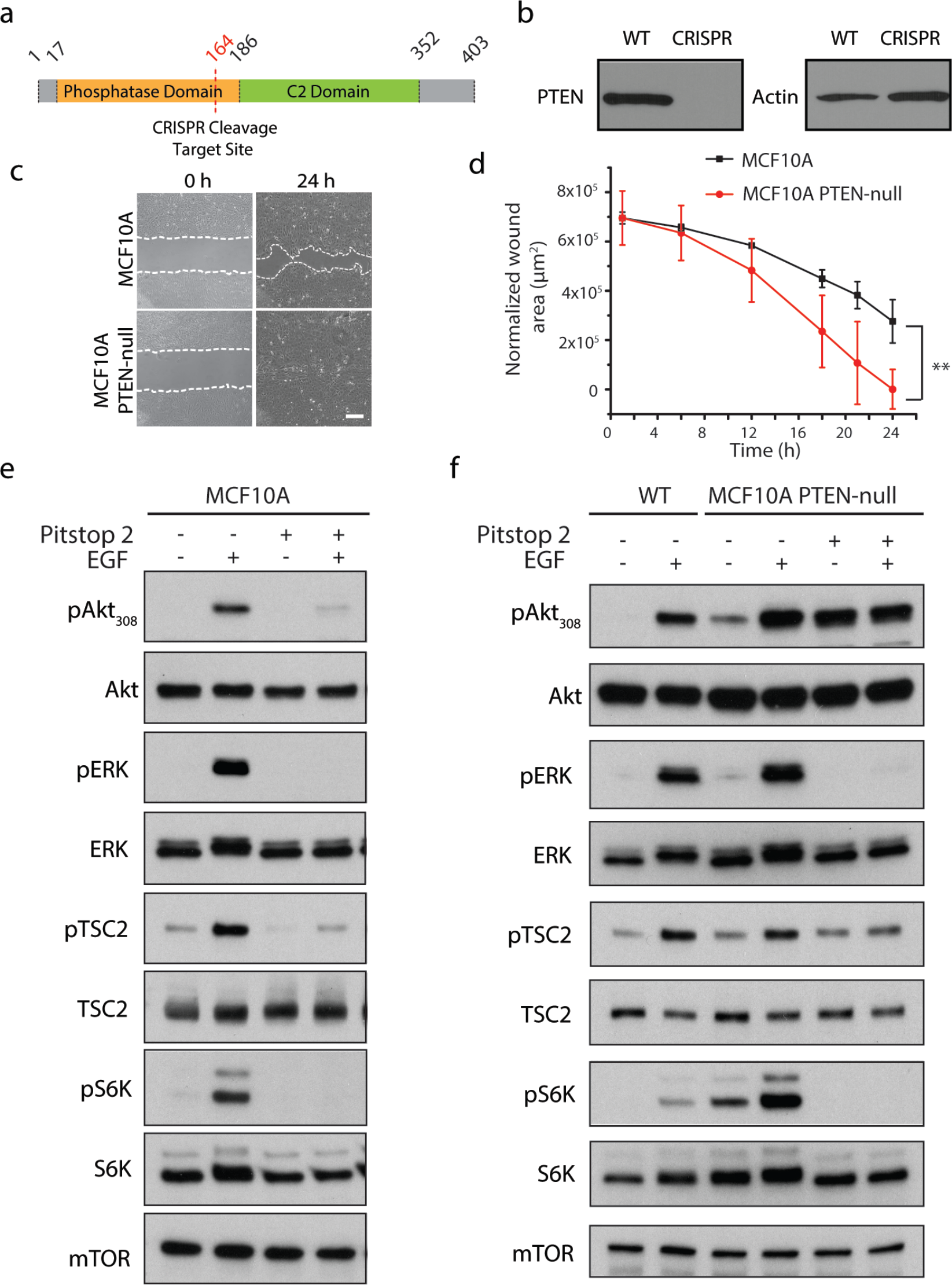
Clathrin-mediated regulation of EGF-induced signaling by PTEN. **a)** Schematic for CRISPR editing of *PTEN* gene. CRISPR cleavage target site was at position 164 in the phosphatase domain of *PTEN* gene. **b)** Western blot showing a complete knockout of PTEN expression in the CRISPR PTEN-null MCF10A cells. **c)** Representative phase contrast images of the wound healing assay at 0 hours and 24 hours for MCF10A and MCF10A PTEN-null cells. White dashed lines mark the boundary between cells and cell-free area. **d)** Quantification of the ‘wound’ area as a function of time. The experiment was performed in quadruplicate (mean ± S.D.). ** *p* < 0.01. **e)** MCF10A cells were treated or not with 20 ng/ml EGF and/or 20 µM Pitstop2 as indicated. Whole-cell lysates were collected after 10 minutes of EGF stimulation and resolved by immunoblotting and probed with the indicated antibodies. **f)** MCF10A and MCF10A PTEN-null cells were treated or not with EGF and/or Pitstop 2 as indicated. Whole-cell lysates were probed with the indicated antibodies. Western blot analysis was performed for three independent experiments.

Next, we used a small-molecule inhibitor of clathrin, Pitstop 2 ^38^, to investigate the requirement of proper clathrin assembly on EGF-induced EGFR activation of the PI3K/Akt/mTORC1 signaling axis. Pitstop 2 inhibits binding of endocytic accessory proteins (such as amphiphysin) to the N-terminal domain of clathrin thereby perturbing proper clathrin network assembly and internalization ^38^. Cells were treated with 20 µM Pitstop 2, which did not impact cell viability, and effectively inhibited EGF internalization (Sup. Fig. 2a, b), as previously reported ^39^. In MCF10A cells, EGF stimulation activated mTORC1 signaling (measured by phosphorylation of pS6K), through PI3K (as seen by increased levels of pAkt_308_) and MAPK/ERK (as seen by increased levels of pERK) (Fig. 1e). Upon Pitstop 2 treatment, EGF is unable to activate Akt_308_ or ERK phosphorylation, thus completely blocking EGF-induced mTORC1 activation. This result suggests that phosphorylation of Akt_308_ and ERK are dependent on clathrin assembly and that clathrin structures may function as a platform for signal transduction ^39^. Genetic inhibition of clathrin using shRNA gene silencing also confirmed the same effect on Akt_308_ observed with Pitstop 2 treatment (Sup. Fig. 2c & d). Interestingly, in PTEN-null cells, which present higher basal levels of pAkt_308_, Pitstop 2 treatment did not reduce Akt_308_ phosphorylation as it did in wild-type cells, but unexpectedly, it increased Akt’s steady-state phosphorylation without EGF stimulation (Fig. 1f). The same effect was observed also at the Ser473 phosphorylation site of Akt (Sup. Fig. 2e), further supporting that Akt activation in PTEN-null cells is indeed linked to clathrin network, and that the reduction of signaling by Pitstop 2 in wild-type cells was not due to cytotoxic effects. However, the clathrin assembly inhibitor still blocked mTORC1 activation, perhaps through its effect on pERK activation. Moreover, EGF induced TSC2 phosphorylation in wild-type and PTEN-null cells (Fig. 1e, f). Similarly, Pitstop 2 treatment blocked EGF-induced TSC2 phosphorylation even though Akt is hyper phosphorylated in PTEN-null cells under the same conditions. Altogether, these results suggest that CME and subsequent vesicle trafficking play a major role in the regulation of the Akt/TSC2 pathway. At the same time, the activation of Akt by Pitstop 2 in PTEN-null cells uncouples Akt phosphorylation from mTORC1 signaling, suggesting that the clathrin platform and vesicle internalization were necessary for EGF activation of mTORC1.

**Figure 2.**
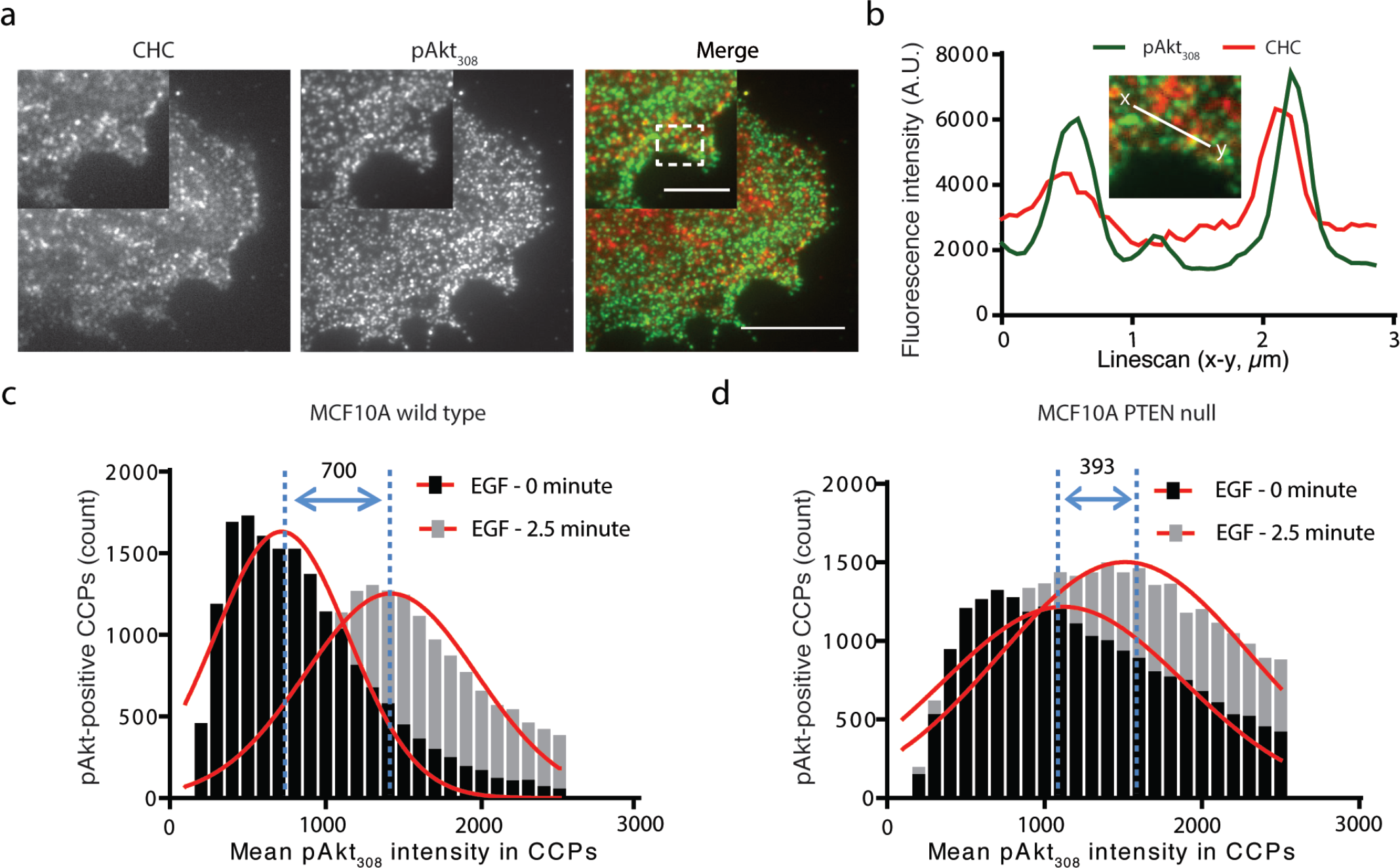
Image quantification of pAKT_308_ levels in CCPs with and without EGF. MCF10A and MCF10A PTEN-null cells treated and not treated with EGF were fixed and immunolabelled cells with CHC and pAKT_308_ antibodies for TIRF imaging. **a)** Representative TIRF image of EGF stimulated MCF10A cell. Insets show high magnification single stained and merged images. Scale bar is 10 µm and 5 µm (inset). **b)** Line scan fluorescence intensity profile (xy) of pAkt_308_ (green) and CHC (red) in a region marked by the white dashed rectangle. **c-d)** Quantification of pAkt_308_ mean intensity in single CCPs was performed as described in the material and methods section. Histogram plot of pAkt_308_ mean intensity in CCPs for MCF10A cells (c) and MCF10A PTEN-null cells (d) with and without EGF. For each condition a total of 10 images was quantified. Bin size is 100. A Gaussian fit for each curve is shown in red. The number above the blue doubled arrow denotes the mean intensity (dashed blue lines) difference between the two Gaussians.

### pAkt level is higher in CCPs in cells treated with EGF

Our Western blot analysis suggested that pAkt is dependent on CME and elevated if endocytosis is blocked in PTEN-null cells. However, this does not necessarily mean that CCPs can act as signaling platforms. We next asked whether pAkt could be enriched in CCPs upon EGF stimulation. In MCF10A cells immunolabeled with clathrin heavy chain (CHC) and pAkt_308_ antibodies (Figure 2a), we measured the fluorescence intensity of pAkt within CCP masks generated from our detection algorithm (as described in materials and methods). The primary goal here is not to claim colocalization, but rather to use pAkt as a measurement of activated Akt within the CCPs. The line scan analysis showed partial overlap between CHC and pAkt puncta (Figure 2b), suggesting that CCPs could indeed contain more pAkt. In both MCF10A wild type (Figure 2c) and PTEN-null (Figure 2d) cells, EGF treatment after 2.5 minutes clearly increased the mean pAkt intensity within CCPs relative to no EGF conditions. Moreover, there was an increased level of pAkt (measured by its mean intensity) in PTEN-null cells (1120) compared with wild type cells (717.5), corroborating our Western results. This result also provides evidence that CCPs can function as signaling platforms.

### PTEN is recruited to small, short-lived CCPs

The analysis of pAkt in CCPs was performed in fixed cells and it is unclear if any of the EGFR signaling machinery might be preferentially localized to CCPs of different lifetimes. Since PTEN-null cells have elevated pAkt signaling, it is plausible that PTEN can attenuate Akt signaling in the proximity of clathrin structures. To investigate if there is an association of PTEN within CCPs, we reconstituted GFP-tagged PTEN (PTEN-GFP) in the SUM149 mCherry-Clc stable cell line via lentiviral transduction (Sup Fig. 3a). To assure that the PTEN-GFP reconstitution was functional, we performed Western blot analysis probing downstream signaling effectors and found that PTEN-GFP expression suppressed basal levels of Akt phosphorylation at Ser473 (Sup. Fig. 3b). As shown in Figure 3a (left panels), there was extensive colocalization of PTEN-GFP with mCherry-Clc in fixed cells. Furthermore, three-dimensional (3D) colocalization was confirmed by 3D reconstruction of these confocal images (Fig. 3a, right panels), in a similar manner to what was done previously ^33^, which further confirms the vesicular association of PTEN with clathrin in these cells.

**Figure 3.**
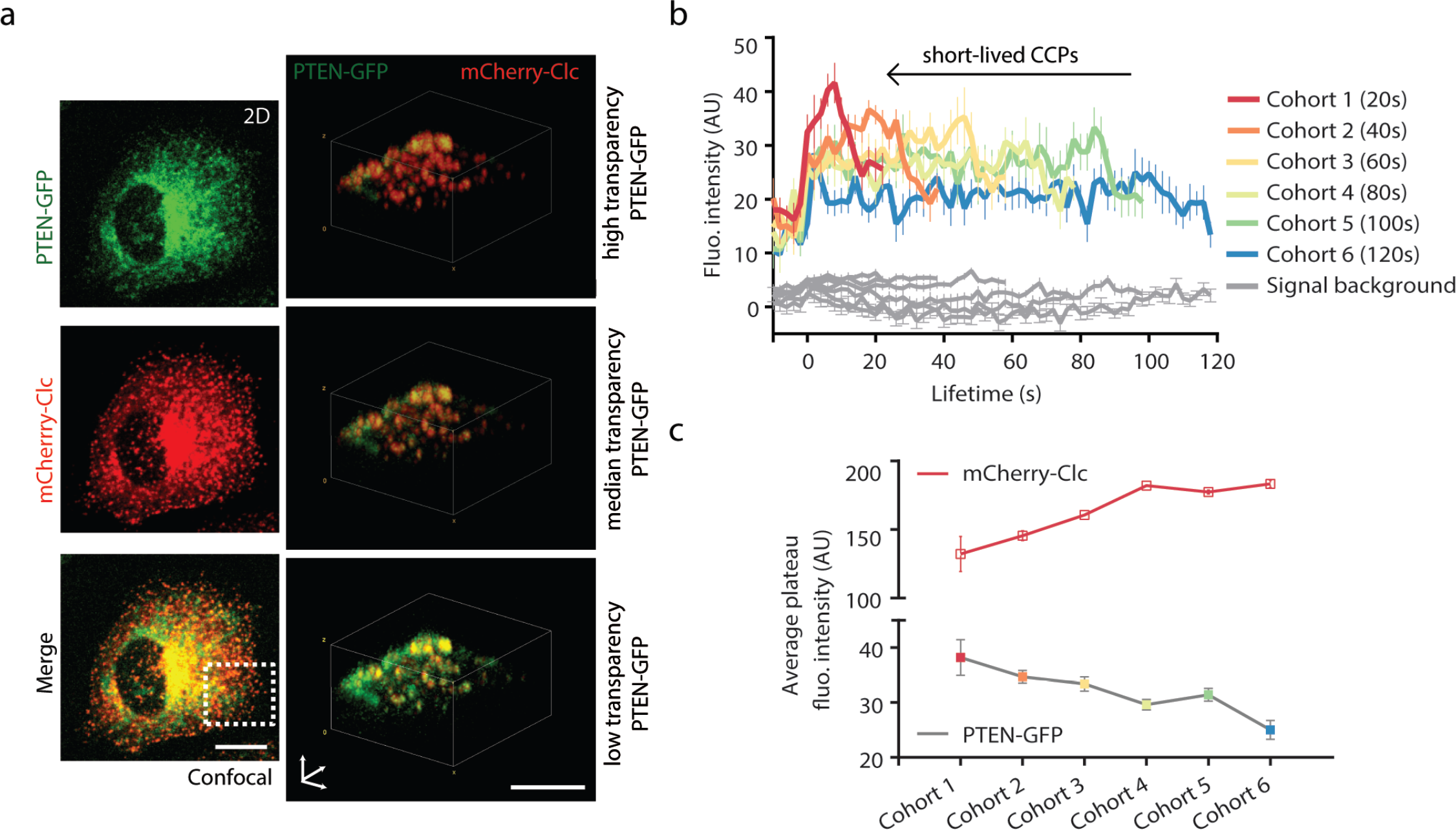
PTEN is preferentially localized to short-lived CCPs. **a)** SUM149 cells stably expressing mCherry-Clc and transduced with PTEN-GFP. Left panels, confocal images show localization of PTEN-GFP (top), mCherry-Clc (middle), and merged channels (bottom) in the 2D image. Right panel shows 3D reconstruction from cell bottom to top of the dash-boxed area closed to the cell edge as marked in the 2D merged image. Three density settings are displayed for PTEN-GFP to allow complete visualization of the signal overlap with mCherry-Clc channel. **b)** Average intensity trajectories of PTEN-GFP in PTEN-positive and PTEN-negative CCPs in 6 lifetime cohorts (20, 40, 60, 80, 100 and 120s) as indicated. Gray traces are PTEN-GFP background signal detected in PTEN-negative CCPs. **c)** Average plateau intensity for PTEN-GFP (colored squares) and CCPs (red squares) for each lifetime cohort. Average plateau intensity was obtained by averaging 5 maximum fluorescent intensity values in each lifetime cohort.

Based upon our observation of PTEN/clathrin colocalization in fixed cells, we sought to characterize this association between PTEN and CCPs using live cell imaging. Recent work has shown that PTEN is tethered to endocytic vesicles in the cytosol ^33^; however, the nature of its localization to the plasma membrane is still not fully understood. To determine whether PTEN association to clathrin endocytic vesicles starts at the plasma membrane, we imaged SUM149 cells stably expressing PTEN-GFP and mCherry-Clc using dual-channel time-lapse TIRF microscopy. In the TIRF field, PTEN-GFP appeared diffuse throughout the plasma membrane while mCherry-labeled CCPs appeared as well-defined puncta in the TIRF field, as expected. The lack of obvious visual colocalization of PTEN with CCPs was not unexpected, as other lipid modifying enzymes have been reported to lack direct association with CCPs at the plasma membrane ^16^. However, with the aid of automated detection and tracking analysis software, we could detect PTEN-GFP signal above background colocalized with a fraction of *bona fide* CCPs. To better characterize the nature of the PTEN association to CCPs, we binned and averaged intensity traces of all CCPs within six lifetime cohorts ranging from 10 – 120 s, similar to what others have done previously ^40^. We then measured PTEN-GFP fluorescence intensity for individual CCPs for each lifetime cohort, as shown in Figure 3b. As measured by its average plateau intensity, the highest PTEN-GFP signal was present in short-lived CCPs (< 20 s) and the plateau intensity gradually decayed as CCP lifetime increased. By comparison, plateau intensity for CCPs increased with longer CCP lifetime cohorts, as we have previously reported ^7^ (Fig. 3c). This result strongly supports that PTEN is preferentially recruited to the subset of small, short-lived CCPs.

### Short-lived CCPs play a vital role in signal transduction pathways

We next asked whether EGFR (hence signaling) could localize to short-lived CCPs, as PI(3,4,5)P_3_ generated by the activation of EGFR is a substrate for PTEN. To analyze EGF/EGFR internalization through CCPs, we stimulated MCF10A cells expressing mCherry-Clc with EGF-Alexa Fluor 647 (AF647) during time-lapse imaging at 37°C. EGF-AF647 addition resulted in rapid binding and clustering of EGFR in CCPs (Fig. 4a). Consistent with what others have observed ^39^, EGF-AF647 colocalized with diffraction-limited CCPs near the cell’s periphery (Fig. 4a). Automated quantitative analysis of mCherry-Clc and EGF-AF647 across multiple cells revealed that 11% of the CCPs showed significant EGF colocalization. CCPs were classified as positive for EGF if the EGF-AF647 signal was significantly higher than the signal for a random association ^41^. Lifetime cohort analysis revealed a preferentially higher degree of association of EGF specific for the < 20 s lifetime cohort (Fig. 4b), which has never been described previously. A late recruitment of EGF in the 80 s and 120 s lifetime cohorts was also observed; however, the plateau fluorescence intensity of EGF-AF647 in CCP lifetime cohorts of 80 and 120 s were about 35% lower than that of the 20 s lifetime cohorts. These data suggest that EGF-occupied EGFR enter CCPs at different kinetic phases and are preferentially associated to short-lived CCPs. Thus, we speculate that these short-lived CCPs are important for signal transduction pathways due to preferential association with the tumor suppressor PTEN and EGF-bound EGFR.

**Figure 4.**
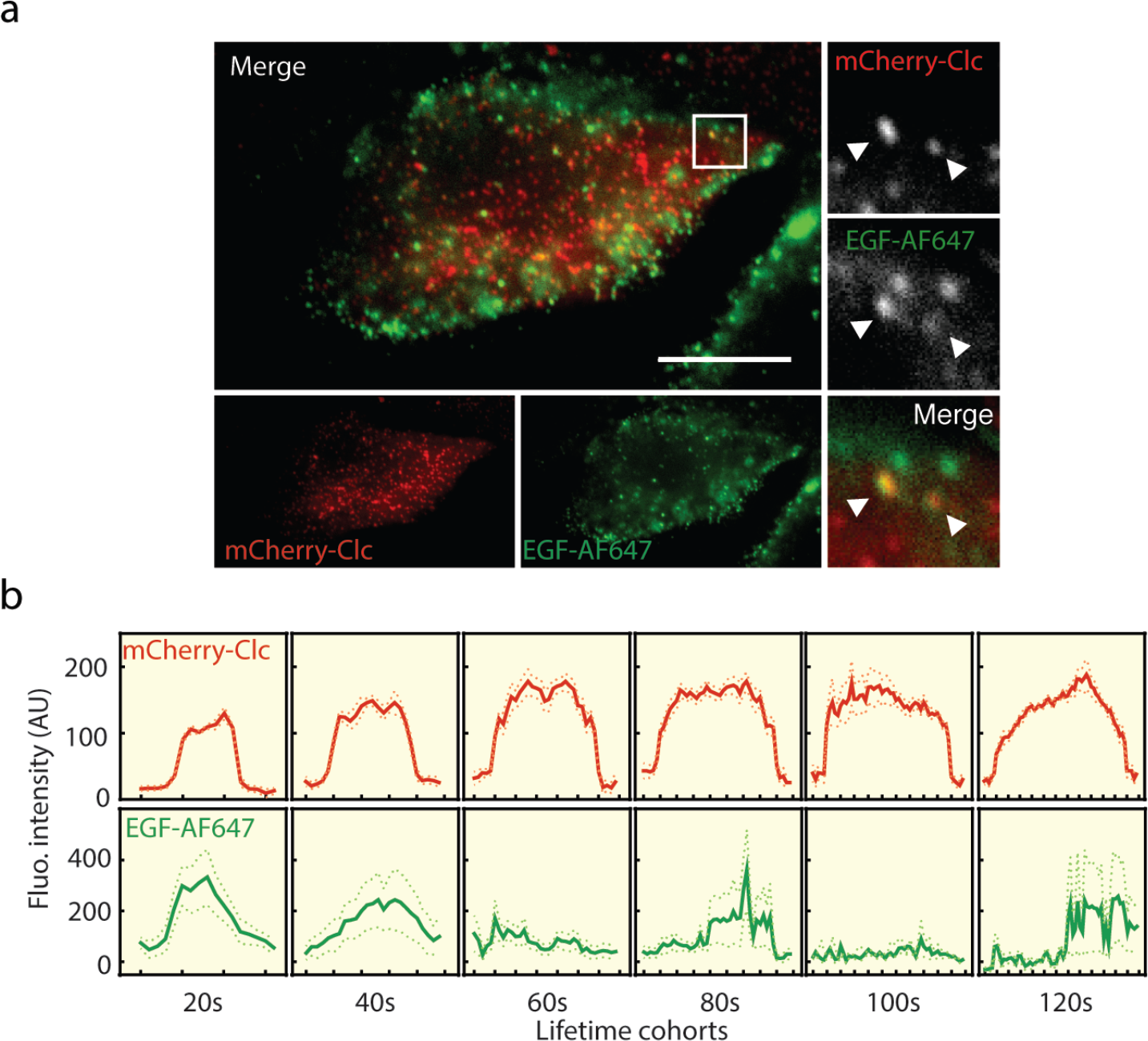
EGF bound EGFR is localized preferentially to short-lived CCPs in MCF10A cells. **a)** Dual-channel time-lapse TIRF imaging of MCF10A cells stably expressing mCherry-Clc and stimulated with EGF-AF647 was performed immediately upon EGF addition at 37°C. Single channels and merged image at initial time point frame after EGF-AF647 stimulation are shown. Insets show high magnification single and merged images of the region marked by the white rectangle. White arrows show examples of colocalization of EGF-AF647 with CCPs. Scale bar is 20 µm. **b)** Average clathrin (top panels) and EGF-AF647 (bottom panels) fluorescence intensity traces in 6 CCP lifetime cohorts. Dotted lines represent SEM.

### PTEN-depleted cells present higher short-lived CCPs than their isogenic counterparts

A growing body of evidence suggests that CME is dysregulated in cancer ^42,43^. Despite this general recognition, it remains unknown how the distinct dynamics of CME that are responsible for the endocytosis of many cargos could be reprogramed in cancer cells. MDA231 and SUM149 cell lines are widely employed, but distinct models of triple negative breast cancer (breast cancer that is estrogen receptor negative, progesterone receptor negative, and lacks HER2 overexpression). SUM149 cells are additionally characterized as an inflammatory breast cancer cell line that lacks endogenous PTEN expression. To gain insight into CCP structures present in these highly aggressive breast cancer cell lines and the normal-like breast epithelial cell line MCF10A, we performed stochastic optical reconstruction microscopy (STORM) on immunolabeled endogenous clathrin heavy chain (CHC) (Fig. 5a). The reconstructed STORM images show substantial resolution improvement over conventional TIRF images. While CCPs appeared as diffraction-limited puncta with no discernible morphological details in TIRF images (Sup. Fig. 4a), in STORM images, CCPs were resolved at a higher resolution with the appearance of ring structures for all three cell lines (Fig. 5a, bottom panel), as others have observed previously ^44^. Notably, we observed larger clusters of clathrin structures in SUM149 cells in some instances (Fig. 5b). To obtain a relative comparison of individual CCP sizes from the STORM images, we computed the radial distribution function *g(r)* from each detected fluorophore ^45^. The *g(r)* curves monotonically decrease with increasing radius. The corresponding correlation length from fitting *g(r)* to an exponential function yields a measure of average CPP radius that can be compared across cell types. SUM149 cells show significantly smaller individual CCPs, on average, when compared to MCF10A and MDA231 cells (Fig. 5c). Although the clusters of clathrin structures were identified only in SUM149 cells, these cells also have smaller CCPs than the MDA231 non-inflammatory triple negative breast cancer cell or MCF10A.

**Figure 5.**
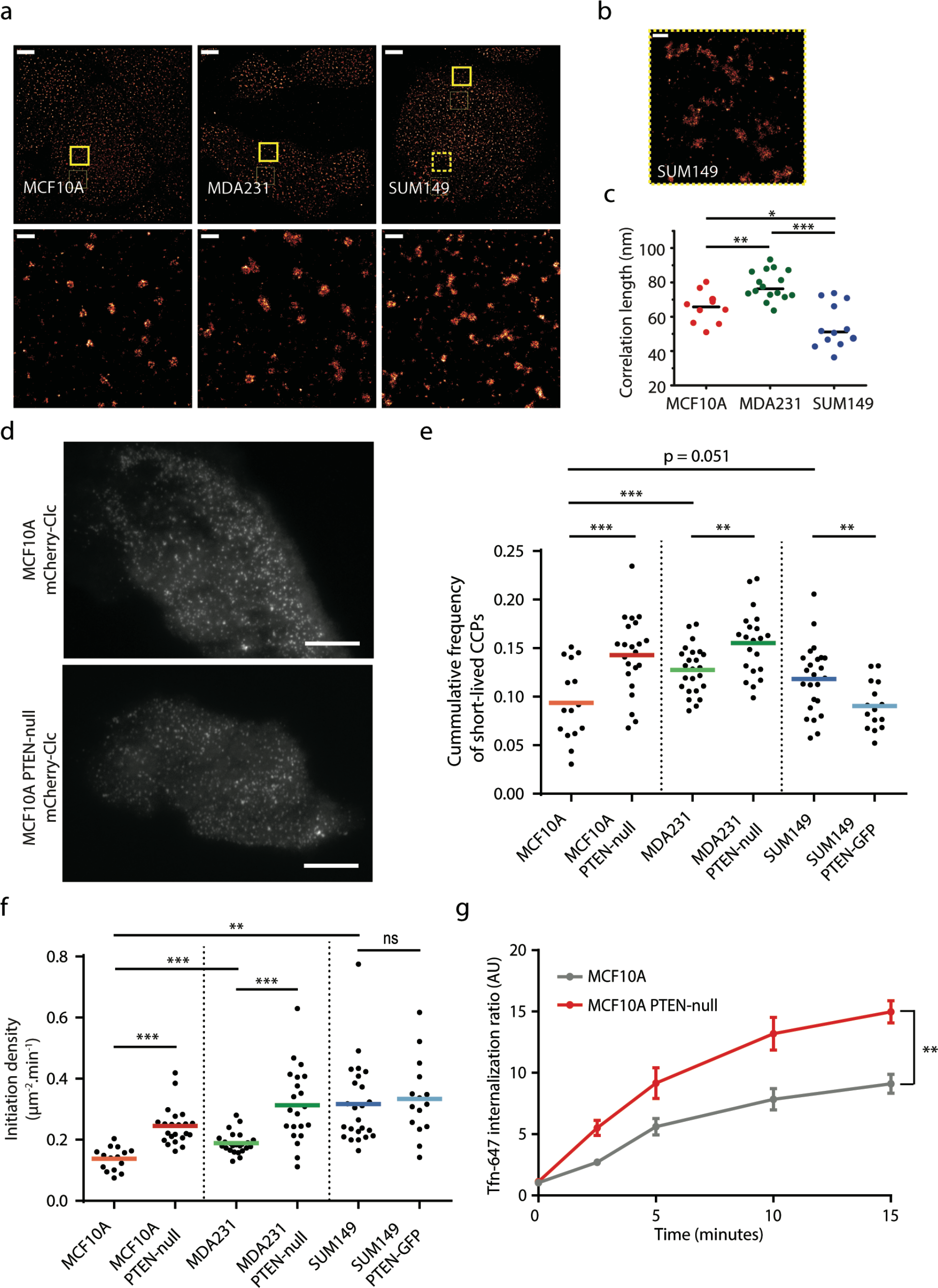
PTEN modulates proportion of short-lived CCPs and initiation density. Super-resolution imaging of CCPs in MCF10A, MDA231, and SUM149 cell lines. **a)** STORM TIRF images of immunostained CCPs in MCF10A, MDA231 and SUM149 cells (upper panel). Zoomed-in areas show solid boxed regions (bottom panel). Images were rendered at 15 nm pixel size. Scale bars are 5 µm and 500 nm for full size and zoomed images, respectively. **b)** Zoomed-in area from dash boxed SUM149 cell image shows a region of the cell with larger clathrin structures **c)** The radial distribution function (*g(r)*) was calculated from single cells within STORM images as a function of probe separation distance (r). Correlation lengths were obtained from fitting g(r) curves for each cell. Quantification of correlation length is shown for MCF10A, MDA231 and SUM149 cells. The solid black line indicates the mean value of 10, 16, and 12 cells analyzed for MCF10A, MDA231, and SUM149, respectively. *p* < 0.05 for all pairs of comparison. * *p* < 0.05, ** *p* < 0.01, *** *p* < 0.001 by two-tailed *t*-test. **d)** Representative TIRF images of CCPs for MCF10A stably expressing mCherry-Clc in comparison to CRISPR PTEN-null MCF10A cells. Scale bar is 10 μm. **e)** Cumulative frequency of short-lived CCPs (10–18 s lifetime) for MCF10A (n = 15), MCF10A PTEN-null (n = 22), MDA231 (n = 23), MDA231 PTEN-null (n = 21), SUM149 (n = 24), and SUM149 PTEN-GFP (n = 15) cells. Each point represents data from a single cell and colored bar represents the mean cumulative frequency for each condition. **f)** Initiation density of CCPs for each of the cell lines as described in b). Each point represents data from a single cell and colored bar represents the mean initiation density for each condition. **g)** Transferrin uptake by MCF10A and MCF10A PTEN-null cells assayed by flow cytometry (n=3, mean ± S.D). ** *p* < 0.01, *** *p* < 0.001 by Wilcoxon test for b) and c) and by paired *t*-test in d).

If short-lived CCPs recruit PTEN and are involved in EGF signaling, then we expect modulation of PTEN level may regulate the fraction of short-lived CCP formation. To test this hypothesis, we generated stable cell lines expressing mCherry-Clc in otherwise isogenic MCF10A and MDA231 PTEN-null cells (Fig. 5d and Sup. Fig. 4b). Using TIRF microscopy, we acquired time-lapse movies of fluorescently labeled CCPs in MCF10A, MCF10A PTEN-null, MDA231 (which have native PTEN expression), MDA231 PTEN-null, SUM149 and, SUM149 + PTEN cell lines. We analyzed only diffraction-limited *bona fide* CCPs, and CCP lifetime distribution and CCP initiation density were obtained by automated tracking and analysis ^41,46^. CCP lifetime represents the length of a trajectory between appearance and disappearance of a fluorescent CCP puncta at the plasma membrane. CCP initiation density is defined as the number of CCPs per unit area per unit time ^3,9,40^.

It is well documented that the heterogeneity of CCP lifetime distribution is in part due to the coexistence of both short- and long-lived CCPs ^21,47^. Interestingly, a higher proportion of short-lived CCPs was observed in MDA231 and SUM149 cells compared to MCF10A cells (Fig. 5e). Additionally, both MDA231 and SUM149 cells have increased rates of CCP initiation density compared to MCF10A (Fig. 5f). These results confirm distinct, cell line dependent CCP dynamics and morphologies. The increase in both short-lived CCPs and CCP initiation densities observed in the cancer cell lines with aberrant signaling compared to the normal cells is consistent with the idea that short-lived CCPs may endow cancer cells with higher signaling capacity. Interestingly, we observed a higher proportion of short-lived CCPs and a higher initiation density in PTEN-null cells (Fig. 5e and f) for both MCF10A and MDA231, a phenotype more similar to SUM149 cells. This finding also agrees well with the correlation between an increase in basal signaling (Fig. 1f) and an increase in short-lived CCPs (Fig. 5e). Consistent with these results, PTEN-GFP reconstitution in SUM149 cells decreased the proportion of short-lived CCPs compared to SUM149 cells (Fig. 5e). However, we did not observe a difference in the initiation density between SUM149 cells and SUM149 cells reconstituted with PTEN-GFP (Fig. 5f). The unaltered initiation rate observed in SUM149 PTEN-GFP reconstituted cells may be a result of long standing signaling adaptations in SUM149 cells that circumvent PTEN necessity. Nevertheless, these data suggest a role for PTEN in regulating CCP lifetime and clathrin-mediated signaling in cancer cells.

There is growing consensus for classifying CCPs with lifetimes below 20 s as abortive CCPs ^4,41,48^. In accordance with this, an increase in short-lived, presumably abortive, CCPs would lead to less cargo uptake. In the case of PTEN-null cells however, an increase in initiation density could offset this decrease in cargo internalization. Interestingly, we found MCF10A PTEN-null cells have significantly higher CME efficiency, as measured by transferrin (Tfn) uptake using Tfn-AF647 as a canonical cargo for CME (Fig. 5g). The nearly two-fold increase in Tfn uptake in PTEN-null cells in part suggests that perhaps not all short-lived CCPs are abortive. Consistent with this, we also found EGF internalization to be higher for MCF10A PTEN-null cells (with higher short-lived CCPs) compared to MCF10A cells at short time points (Sup. Fig. 2b).

### PI(3,4,5)P_3_/AM increases short-lived CCPs

A direct effect of the loss of PTEN is an excessive accumulation of the secondary messenger lipid PI(3,4,5)P_3_ at the plasma membrane ^49^. To test whether increased PI(3,4,5)P_3_ concentrations typically found in some advanced types of breast cancer ^50^ would directly affect CCP dynamics, we used a synthetic membrane-permeant phosphoinositide derivative, PI(3,4,5)P_3_/AM (acetoxymethyl ester) to acutely elevate PI(3,4,5)P_3_ concentration at the plasma membrane ^51^ (Fig. 6a). Serum-starved MCF10A mCherry-Clc cells were treated with PI(3,4,5)P_3_/AM followed by time-lapse live cell TIRF imaging and analysis. Intriguingly, PI(3,4,5)P_3_/AM treatment resulted in a significant increase in the proportion of short-lived CCPs (Fig. 6b), mirroring the effect in PTEN-null cell lines (Fig. 5e). In contrast, we found no significant difference in the fraction of short-lived CCP population upon EGF addition, suggesting that EGF-induced PI3K activation and subsequent local increase in the PI(3,4,5)P_3_ pool is not sufficient to alter CCP lifetime distribution (Fig. 6b). Consistent with this, we also did not observe a change in the proportion of short-lived CCPs in PI(3,4,5)P_3_/AM-treated cells when stimulated with EGF, suggesting that global concentration changes in PI(3,4,5)P_3_, similar to the case when PTEN is deleted, directly mediate changes in CCP dynamics and signaling.

**Figure 6.**
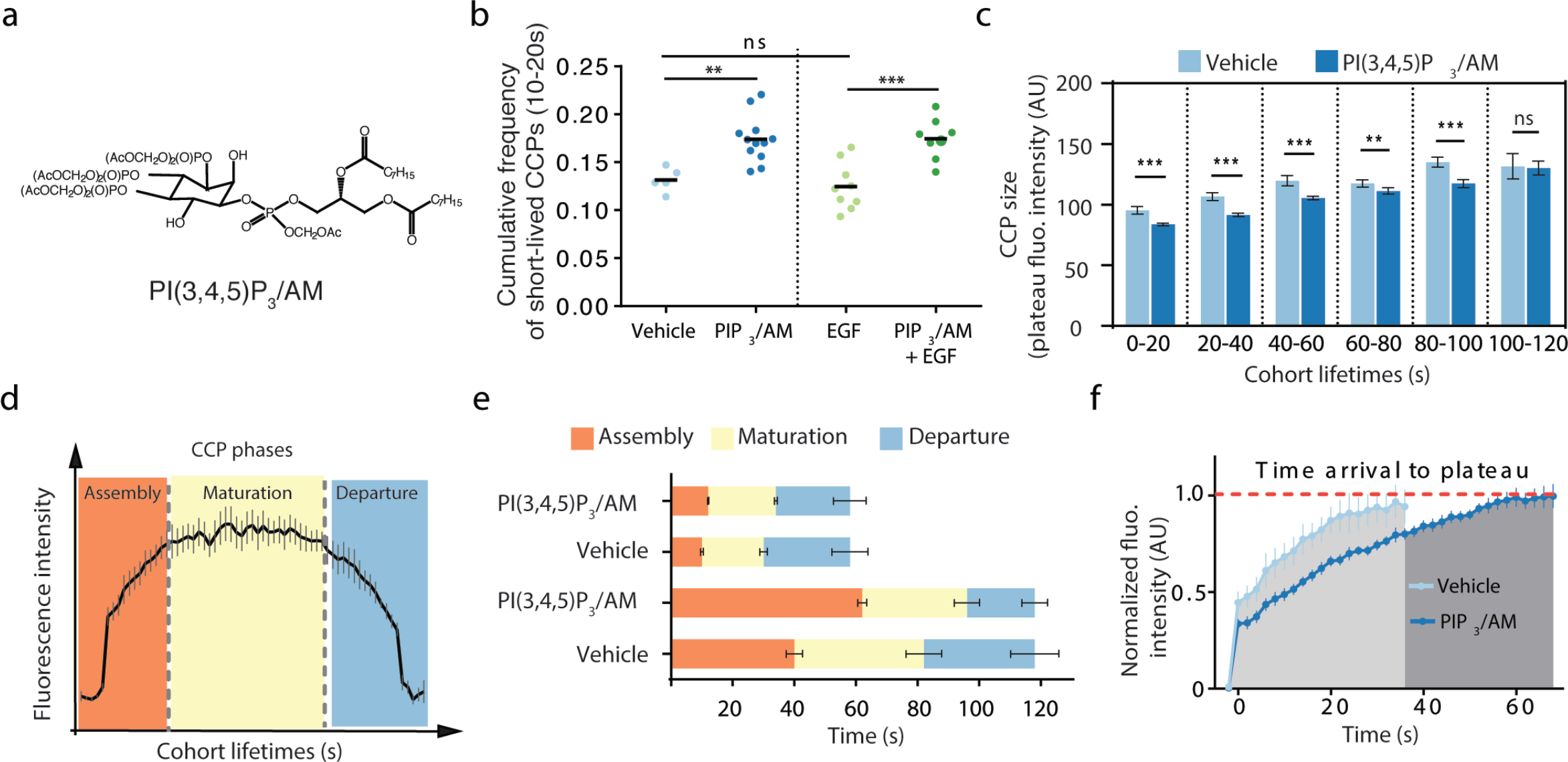
Membrane permeant PI(3,4,5)P_3_/AM changes kinetics of CCP assembly. **a)** PI(3,4,5)P_3_/AM chemical structure. **b)** Cumulative frequency of short-lived CCPs (10–18 s lifetime) in MCF10A mCherry-Clc cells treated with vehicle (DMSO, n=5), PI(3,4,5)P_3_/AM (n=12), EGF (n=9), or PI(3,4,5)P_3_/AM plus EGF (n=8). **c)** Average plateau intensity of all six lifetime cohorts (20, 40, 60, 80, 100, and 120 s) for vehicle and PI(3,4,5)P_3_/AM treatment in MCF10A mCherry-Clc cells. Error bars represent SE. **d)** Schematic showing a fluorescence intensity trace for a CCP lifetime cohort. The three CCP phases (assembly, maturation and departure) are marked to correspond to regions of clathrin intensity growth, plateau, and decay. **e)** The duration of CCP phases averaged over all extracted trajectories from the 100–120 s lifetime cohort for PI(3,4,5)P_3_/AM treatment and vehicle conditions. Error bars represent SE. **f)** Comparison of intensity traces for the 100–120 s cohort between vehicle and PI(3,4,5)P_3_/AM conditions. The arrival to plateau time point was determined by the first time point that the fluorescence intensity to reach 95% of the maximum fluorescence intensity. The average slopes of the fluorescence increase were 0.018 ± 0.002 and 0.01 ± 0.0006 AU/s for vehicle and PI(3,4,5)P_3_/AM conditions, respectively. ** *p* < 0.01, *** *p* < 0.001 by Wilcoxon test and *t*-test for b) and c), respectively.

Previous study has shown that an increase in PI(4,5)P_2_ by overexpressing PIP5Kα increases CCP size ^16^. Our super-resolution imaging revealed smaller CCP sizes in PTEN-deleted SUM149 cells (Fig. 5a & c). Thus, we investigated whether PI(3,4,5)P_3_ might also have an effect on CCP size. By quantifying plateau intensity over the lifetime of CCPs as a global measure of size ^9,16^, we found a reduction in clathrin plateau intensity in PI(3,4,5)P_3_/AM-treated cells across all the lifetime cohorts except for the 100–120 s cohort (Fig. 6c). The decrease in CCP size across most of the cohorts is consistent with our observed smaller CCPs in SUM149 cells. The increase, plateau (defined as 95% of the maximum averaged intensity), and the sudden decrease in intensity tracks of lifetime cohorts corresponding to the growth, maturation, and scission or disassembly of CCPs, respectively (Fig. 6d) ^40,52^. Even though PI(3,4,5)P_3_ does not alter the plateau intensity for the 100–120 s cohort, it remodels the distinct phases of the CCP life cycle. Specifically, for long-lived CCPs, PI(3,4,5)P_3_ prolongs the assembly phase at the expense of maturation and departure (Fig. 6e). In addition to extending the assembly phase, PI(3,4,5)P_3_ also reduced the clathrin assembly rate from 0.018 ± 0.002 to 0.01 ± 0.0006 AU/s (Fig. 6f). Together, these results provide strong support that PTEN’s regulation on CCP dynamics is directly mediated by its function in regulating PI(3,4,5)P_3_ at the cell membrane.

## Discussion

The paradigm of functional specialization of CCPs has challenged the conventional view that CCPs are generally a uniform population of endocytic machineries where different cargo molecules are passive passengers in the endocytic itinerary and they do not influence dynamics by their very nature. Instead, an increasing body of work has revealed that both signaling receptor like GPCRs or EGFRs and nutrient receptors like TfnRs exert cargo-dependent control on CCP composition or dynamics ^5,7,39,53^. GPCR-mediated delay of CCP dynamics was thought to delay receptor resensitization by increasing the time of GPCR-arrestin interaction at the plasma membrane ^5^. Instead of a kinetic mechanism to generate functional specialization, we have identified a ‘signaling-capable’ functional specialization of CCPs (Fig. 7). An earlier study suggested that EGFR signaling that leads to Akt phosphorylation is regulated by a partially distinct CCP population ^39^. Here we detected a major presence of ligand-stimulated EGFR in the same CCP cohorts (up to 40 s) where PTEN was also enriched. These signaling-capable CCPs are short-lived and can be promoted by an increase in PI (3,4,5)P_3_ level, both through an acute increase in plasma membrane PI (3,4,5)P_3_ level or through deletion of PTEN. The intriguing connection between elevated basal PI3K signaling and morphological CCP phenotypes in PTEN-deleted SUM149 cells support the existence of signaling-capable short-lived CCPs. Interestingly, EGF-induced signaling depends on clathrin, as others have shown as well via various clathrin inhibition approaches ^39^, supporting the model of a clathrin scaffold acting as a signaling platform. However, it was surprising and interesting that Pitstop 2 that blocks clathrin interaction with accessory proteins led to higher Akt phosphorylation without EGF stimulation in MCF10A PTEN-null cells. We speculate that a signaling protein inhibitor is recruited to the plasma membrane clathrin platform in a clathrin- and PI (3,4,5)P_3_-dependent manner. Perhaps more interesting is that both ERK and mTORC1 pathways were blocked when Akt was highly phosphorylated in PTEN-null cells treated with Pitstop 2, pointing to the requirement of vesicle internalization for ERK and mTORC1 signaling. Recent findings that persistent G protein signaling from internalized GPCRs ^54^, active EGFRs remaining at the plasma membrane responsible for continuous ERK signaling ^55^, and macropinosome-dependent mTORC1 signaling in macrophages ^56^ all agree with our proposition of such compartmentalized signaling.

**Figure 7.**
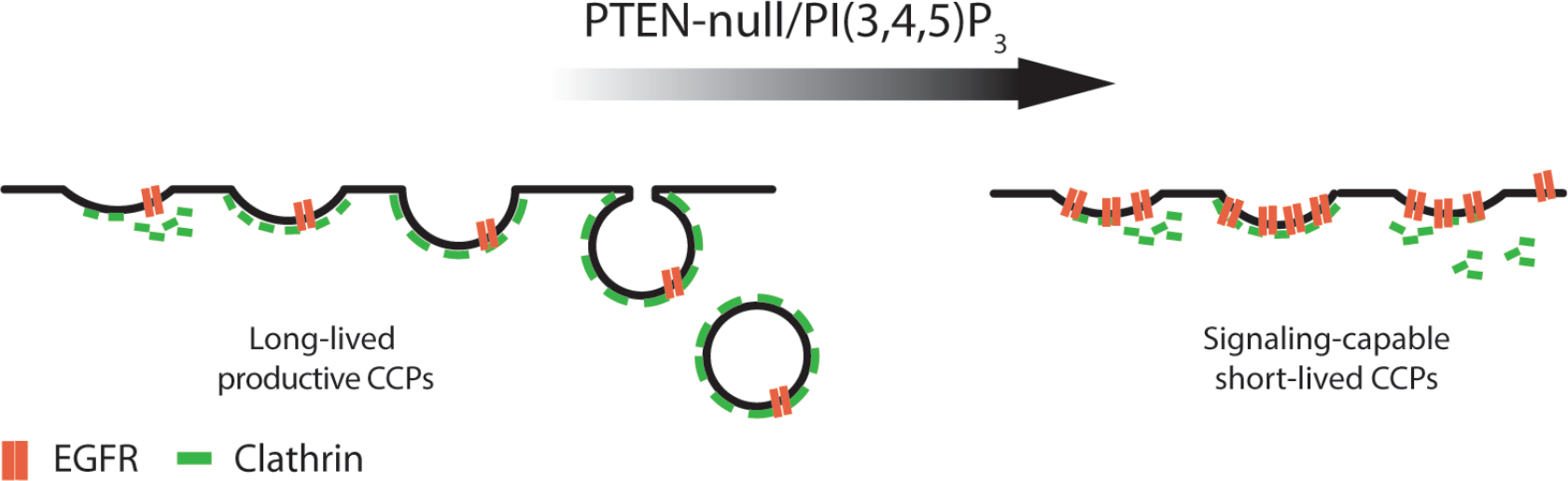
Schematic of model for signaling-capable CCPs. Deletion of PTEN or increase in PI(3,4,5)P_3_ both increase the proportion of short-lived signaling-capable CCPs relative to long-lived, fully internalized CCPs.

The notion of short-lived signaling-capable CCPs appears to contradict the apparently prevailing understanding that CCPs under 20 seconds lifetime are deemed abortive ^4,21,41,48^. These previous studies have alluded to the idea that cargo stabilization and dynamin recruitment govern the transition from abortive to productive CCPs. Interestingly, these studies were performed under no ligand stimulation. In our case, we clearly observed an enrichment of EGFR in short-lived CCPs during EGF stimulation, contrary to the enrichment of TfnR found in long-lived CCPs ^7^. Thus, one could postulate that short-lived CCPs may not represent truly abortive CCPs, at least not in the sense of their signaling function. The overall increase in Tfn and EGF CME in PTEN-null cells where we find a significantly higher proportion of short-lived CCPs also provides evidence that these short-lived CCPs may not be the classically defined abortive CCPs. Whether these short-lived CCPs do internalize or not will be an interesting question to examine in the future in this and other ligand-stimulated receptor systems.

Increased levels of PI(3,4,5)P_3_, which can be accomplished by PTEN deletion, were found to have a direct effect on regulating signaling-capable short-lived CCPs, highlighting a previously unknown function of PI(3,4,5)P_3_. An increase in PI(3,4,5)P_3_ could also lead to an increase in PI(3,4)P_2_ through the action of 5’ phosphatases OCRL and/or SHIP1/2. Both ORCL and SHIP2 are found in CCPs and govern their dynamics ^57,58^. Interestingly, SHIP knockdown, which effectively leads to more PI(3,4,5)P_3_, was found to reduce CCP lifetime ^58^, consistent with what we have found in our study. Moreover, increased level of PI(3,4)P_2_, through deletion of PI3K C2α, was found to attenuate CCP dynamics ^20^. Thus, it is possible that the effect of PI(3,4,5)P_3_ may be through PI(3,4)P_2_ in PTEN-null cells or during acute increase of PI(3,4,5)P_3_. Although PI(3,4,5)P_3_ is converted by PTEN to PI(4,5)P_2_, we do not think the effect of PTEN on CCP dynamics is directly related to changes in the PI(4,5)P_2_ level, since PI(3,4,5)P_3_ is only a small fraction compared to the bulk PI(4,5)P_2_ level^13^. We also do not think the effect of PTEN on CCP dynamics is due to its protein phosphatase activity that has been reported recently to play a role in late endocytic trafficking by dephosphorylating Rab7 ^59^. Instead, we believe PTEN’s role in regulating signaling-capable CCPs is through PI(3,4,5)P_3_ as the lipid mediator, as supported by the same endocytic phenotypes (increased short-lived CCPs and increased initiation density) found in PI(3,4,5)P_3_/AM-treated cells. Interestingly, the increase in short-lived CCPs was not observed in cells stimulated with EGF, suggesting that transient formation of PI(3,4,5)P_3_ due to PI3K activation does not robustly affect CCP dynamics. In agreement with this, a previous study showed that wortmannin, a PI3K inhibitor that causes reduced PI(3,4,5)P_3_ levels, had little effect on transferrin internalization ^60^.

If EGFR is present at high levels in signaling-capable CCPs, why are they short-lived? We find that EGF binds preferentially near the edge of a cell, which is similar to what others have seen ^39^. This is not due to inaccessibility of EGF to the bottom of the cells, as much larger Tfn can easily access the basal cell surface ^21^. We have previously found that increase in cell tension due to cell spreading increases the proportion of short-lived CCPs and initiation density ^9^. It is possible that dense actin networks and focal adhesions typically found at the cell periphery might also provide spatial organization to EGF-induced signaling-capable CCPs, which in turn regulate their lifetime. However, because we also observed higher PTEN localization to short-lived signaling-capable CCPs, it is also possible that PTEN recruitment, through unknown mechanisms, led to shorter CCP lifetimes. Regardless of the mechanism of short-lived CCPs, our results suggest that internalization of signaling receptors through CCPs could be spatially organized to elicit signaling in different membrane domains and possibly different cytoplasmic routes thereafter. Whether or not this phenomenon might depend on cytoskeletal organization requires further investigation.

In a broader sense, the distinct phenotypes and dynamics of the highly aggressive PTEN-deleted SUM149 cancer cell line reinforces the idea that CME is reprogramed in cancer cells ^42,43,61^. It has recently been discovered that many different cancers are caused by mutations in genes that directly or indirectly regulate PI(3,4,5)P_3_ levels in cells. Mutations that cause increased production of PI(3,4,5)P_3_ without a need for receptor stimulation are more likely to result in carcinogenesis. Aberrant cancer cell signaling could be in part due to the increased CCP initiation densities observed in cancer cells, which could in turn, be either a consequence of their active metabolism or their up-regulation of many signaling receptors. More importantly, the loss of PTEN, the major attenuator of the PI3K signaling, directly results in more signaling-capable CCPs. The novel subcellular localization of PTEN to nascent CCPs uncovers a novel biological function of PTEN in controlling CCP dynamics and helps shed light on cancer cell signaling. A deeper understanding of PTEN’s compartmentalized-functions will help guide rational design of novel therapies against new vulnerabilities of cancer cells revealed by CCP dynamics.

## Methods

### Cell culture

MCF10A cells were cultured in DMEM/F12 media (Corning) supplemented with 5% horse serum (HS), 10 μg/mL insulin, 0.5 μg/mL hydrocortisone, 0.02 μg/mL epidermal growth factor, 0.1 μg/mL cholera toxin, 100 U/ml penicillin, 100 μg/ml streptomycin, 2.5 µg/ml amphotericin B, and 5 µg/ml gentamicin. MDA231 cells were maintained in RPMI1640 media (Gibco) supplemented with 10% fetal bovine serum (FBS), 100 U/ml penicillin, 100 μg/ml streptomycin, 2.5 µg/ml amphotericin B, and 5 µg/ml gentamicin. SUM149 cells were maintained in F12 media (Gibco) supplemented with 5% FBS, 5μg/mL insulin, and 1 μg/mL hydrocortisone, 50 U/ml penicillin, 50 μg/ml streptomycin, 2.5 µg/ml amphotericin B, and 5 µg/ml gentamicin. MCF10A and MDA231 cells were grown in 5% CO_2_. SUM149 cells were grown in 10% CO_2_.

### Stable cell lines

MCF10A, MDA231, and SUM149 cell lines were infected with mCherry-Clc lentivirus produced from the pMIEG-mCherry-Clc vector construct (gift from Dr. Sandra Schmid). SUM149 cells were infected with EGFP lentivirus produced from pSMPVW-EGFP (gift from Dr. Andrew Tai) or PTEN-EGFP lentivirus produced from pHR-SIN-PTEN-GFP (gift from Dr. Miho Iijima). Lentiviruses were generated by the University of Michigan Vector Core. Infections were performed in OptiMEM (Gibco) with 10 µg/mL polybrene and 1X or 1.67X virus. Transduced cells were sorted and pooled based on relative fluorescence intensities. CCPs of each pool were imaged and appropriate pool was selected and expanded for further studies.

### PTEN CRISPR cell lines

MCF10A and MDA231 cell lines were transfected using the Nucleofector II system (Lonza) with pSpCas9(BB)-2A-GFP (PX458) ^20^, which was a gift from Dr. Feng Zhang (Addgene plasmid # 48138), containing the target sequence CCAGGGAGTAACTATTCCCA against PTEN. Two days after transfection, single cells were sorted for GFP expression into 96 well plates. Following clonal expansion, genomic DNA was isolated and clones were screened for PTEN mutations using SURVEYOR reactions (IDT) with the following primer pair: Forward-GCTACGACCCAGTTACCATAGC and Reverse-GCCACGTCTTATCACTTCTTCC. Positive clones were sequenced to identify specific mutational events and Western blotted for PTEN (9559; Cell Signaling).

### Immunostaining

Cells were plated onto MatTek dishes and incubated with regular media at 37 °C and 5% CO_2_ overnight. Growth media was then removed and coverslips were washed with phosphate buffer saline (PBS). Cells were permeabilized with 0.05% Triton X-100 in 2% paraformaldehyde (PFA) for 5 minutes followed by fixation with 4% PFA for 30 minutes. Dishes were washed three times with PBS and blocked with 1% bovine serum albumin (BSA) for overnight at 4°C. Primary antibody against clathrin heavy chain (ab2731, Abcam) was incubated for 1 h at room temperature followed by washes and secondary antibody incubation Alexa Fluor 647 (A-21235, Thermo Fisher) for 1 h. Cells were washed three times with PBS and kept in PBS for super-resolution imaging.

### Super-resolution imaging and analysis

Cells were imaged in buffer containing 50 mM Tris, 10 mM NaCl, 10% w/v glucose, 50 mM β-mercaptoethanol, 40 μg mL^-1^ catalase (Sigma), 500 µg mL^-1^ glucose oxidase (Sigma), pH 8.0. Imaging was performed under total internal reflection using an Olympus 1X81-XDC inverted microscope with a cell TIRF module, a 100X APO objective (NA = 1.49), and active Z-drift correction (ZDC) (Olympus America, Center Valley, PA), and a 1.6X magnification in the emission channel. Excitation of Alexa-Fluor 647 was accomplished using a 647 nm diode laser (OBIS 647 LX-100FP, Coherent). Excitation and emission were filtered using the quadband filter cube set ET-405/488/561/647 (Chroma, Bellows Falls, VT) and images were acquired on an iXon Ultra EMCCD camera (Andor Nanotechnology, South Windsor, CT). At least 5000 individual image frames were collected for each reconstructed image.

Single molecules were identified and localized in individual image frames, then processed further to remove outliers, correct for stage drift, tabulate resolution, and reconstruct images using home-built software written in Matlab described previously ^45^. Radial distribution functions, *g*(*r*), were tabulated from single cells within a user-defined region of interest, and *g*(*r*) from single cell images were fit to an exponential function, *g*(*r*)=1+exp(-*r*/*r*_o_) in which *r*_o_ estimates the average CCP radius ^45^. Curves were also fit to a more complicated function that accounts for probe over-counting ^45^, and findings were similar within error.

### Membrane permeant PI(3,4,5)P_3_/AM treatment

PI(3,4,5)P_3_/AM was dissolved in freshly opened dimethyl sulfoxide (DMSO) to a stock solution of 50 mM. The portion to be used was mixed with 10% pluronic F127 in DMSO in 1:1 (v/v) ratio to avoid precipitation when mixed with cell medium. Cell medium was then added to the phosphoinositide to a final concentration of 50 µM PI(3,4,5)P_3_/AM. This solution was directly added to the cells. Individual cells were imaged using time-lapse TIRF microscopy followed by CCP dynamics analysis after 20 minutes of incubation time.

### Cell treatments and Western blotting

For signaling studies, cells were plated in a 6-well plate and allowed to adhere for 24 hours. Cells were serum-starved for 24 hours before being treated with 25 µM Pitstop 2 (Abcam) for 30 minutes and/or 20 ng/ml human EGF (324831; Millipore) for 10 minutes as indicated. Cells were washed in PBS on ice and lysed for 10 minutes in ice cold lysis buffer (40 mM HEPES, pH 7.5, 120 mM NaCl, 1 mM EDTA, 10 mM pyrophosphate, 10 mM glycerolphosphate, 50 mM NaF, 1.5 mM Na_3_VO_4_, 0.3% CHAPS, and a mix of protease inhibitors). Lysates were centrifuged at 12,000 x g for 15 minutes at 4°C, and the supernatant was mixed with 4x SDS sample buffer and denatured for 5 minutes at 95°C. The samples were resolved by Tris-glycine SDS-PAGE and transferred to a nitrocellulose membrane. The membrane was washed; blocked; and probed using the following antibodies: PTEN (9552; Cell Signaling); pAkt_308_ (2965 or 4056; Cell Signaling); pAkt_473_ (4060; Cell Signaling); total Akt (4691, Cell Signaling); pERK (9106 or 4376; Cell Signaling); ERK (4695; Cell Signaling); pS6K (9234; Cell Signaling); S6K (2708; Cell Signaling); pTSC2 (3617; Cell Signaling), TSC2 (4308; Cell Signaling); pRaptor (2083; Cell Signaling); mTOR (2983; Cell Signaling); GAPDH (24778; Santa Cruz); Actin (8432; Santa Cruz); and GFP (290; Abcam). At least three independent experiments were performed to confirm the obtained results.

### Wound healing

Wound healing assays were performed similarly to previously reported work ^62^. Wild type and PTEN-null cells were plated at high confluence in two chambers of the 2-well silicone-culture insert (Ibidi). The insert was removed after 12 hours and the cell-free gap (wound area) was imaged for 48 hours using an Olympus-IX81 spinning disk confocal microscope (CSU-X1; Yokogawa, Japan) coupled with a stage top incubator (PPZI; Tokai Hit, Japan) at 37°C and 5% CO_2_. Quantitative analysis of the open areas was performed using the wound healing pipeline of CellProfiler software (http://cellprofiler.org/) at the indicated time points.

### Transferrin/EGF uptake assay by flow cytometry

For analysis of transferrin or EGF internalization, a 10-cm dish with cells about 70% confluent was incubated for 45 min at 37°C in serum-free medium. Cells were detached from the dish using PBS/5mM EDTA and pelleted at 1,000 rpm for 10 minutes before resuspending in PBS4+ (PBS supplemented with 0.2 mM CaCl_2_, 1 mg/ml BSA, 1 mM EDTA, and 1 mM MgCl_2_). Cells were first incubated on ice for 30 min followed by addition of 5 μg/ml transferrin Alexa Fluor 647 or 20 ng/ml EGF-Alexa Fluor 647 (Thermo Fisher Scientific). Cells were transferred from ice to a 37°C water bath for the indicated amount of time, while keeping one sample on ice for the measurement of total cell surface binding. Uptake was halted by returning cells to ice and 500 ul fresh, ice-cold 0.11% of pronase solution in PBS was added to the samples (excluding the “total” sample) to remove surface-bound ligand for 10 minutes. Cells were pelleted for 1 minute at 10,000 rpm and the pellets were resuspended in 200 µl PBS, followed by addition of 200 µl of 4% paraformaldehyde. Samples were analyzed within 1 hour using a Guava flow cytometer (Millipore). The fluorescence intensity median of each condition was normalized by the total surface binding sample and the data was plotted as internalization ratio over time.

### Total internal reflection fluorescence (TIRF) microscopy and analysis

Cells were prepared for TIRF microscopy as previously described ^7,9^. Briefly, the day before transfection, mCherry-Clc expressing cells were seeded at 60% confluence per well in a six-well plate containing 22 x 22 mm fibronectin (5 µg/ml) coated glass coverslips. The coverslip containing cells were mounted in a home-made imaging chamber and sealed with VALAP (1:1:1 of Vaseline, lanolin, and paraffin). For imaging, the coverslip chamber containing cells were rapidly transferred to a 37°C pre-warmed microscope stage and three to five cells were imaged per coverslip. TIRF imaging was performed using a Nikon TiE-Perfect Focus System (PFS) microscope equipped with an Apochrmomat 100X objective (NA 1.49), a sCMOS camera (Flash 4.0; Hamamatsu Photonics), and a laser launch controlled by an acousto-optical tunable filter (AOTF). Image acquisition was controlled by ImageJ Micro-manager software (NIH). Dual-channel (561nm and 488nm), time-lapse image series were acquired by sequential, nearly simultaneous acquisition of both channels, using 100 to 300 ms exposure at 2 s interval for 10 minutes. All movies of individual cells were acquired using the same TIRF angle for any given pair of control and experimental groups and were acquired on the same day. For each condition, at least 10 movies of different cells were recorded. Fluorescent particle detection, lifetime tracking, and lifetime analysis of CCPs in the TIRF movies were performed using custom-written software in Matlab (MathWorks) as previously described and validated ^3,21,46,9^. Intensity profile analysis on CCP lifetime cohorts was performed as previously described following normalization to the background fluorescence ^6,40^. The frequency of CCPs at each lifetime from 10 to 600 s for each condition was computed to obtain the lifetime distribution, but only data from 10 to 120 s lifetime cohorts were analyzed. Initiation density was computed as described previously by dividing the number of bona fide CCP appearance per unit area and time for each cell ^63^.

For pAkt_308_ quantification in CCPs, 24h serum-starved cells were treated with 5 ng/ml EGF for 2.5 minutes. The cells were fixed in 4% PFA and permeabilized with 0.05% triton-X100 as described above (*Immunostaining* sub-section). Cells were incubated with clathrin heavy chain (ab2731, Abcam) and pAkt_308_ (Abcam) antibodies for 1 h at room temperature followed by washes and secondary antibodies incubation with Alexa Fluor 647 (A-21235, Thermo Fisher) and Alexa Fluor 546 (A-10040, Thermo Fisher), respectively, for 1 h. Dual-channel images of multiple cells were taken by TIRF microscopy for both channels. Using Cme-analysis software in Matlab, we detected and generated masks for CHC and pAkt_308_ images. By multiplying the masks, we extrapolated pAkt_308_ mean intensity in pAkt_308_-positive CCPs. All mean intensity values of each condition were plotted as a histogram and a Gaussian function was used to fit the data using Prism Graph pad software.

## Statistical analysi

For the cumulative frequency of lifetime distribution (from 10 s to 18 s) and initiation density data, Wilcoxon rank sum test was performed. Statistical difference of other data sets was determined by two-tailed Student’s *t* test: \**p* < 0.05, \*\**p* < 0.01, *** *p* < 0.001.

## Acknowledgements

We thank Dr. Miho Iijima (Johns Hopkins University) for the PTEN-GFP construct and Dr. Cartsen Schultz (EMBL) for the lipid permeant PI(3,4,5)P_3_/AM. This work is supported in part by MCubed (to S.M. and A.P.L.), University of Michigan and Pardee Foundation (A.P.L.), the Breast Cancer Research Foundation (SDM), and the Metavivor Foundation (SDM), and National Institutes of Health (GM-110215; J.S.). We acknowledge UM Vector Core (P30 CA046592), UM Flow Cytometry Core, and UM DNA sequencing core - support via the University of Michigan Cancer Center Support Grant (P30 CA046592).

## Author Contributions

L.K.R-M and A.P.L have full access to the data in the study and take responsibility for the integrity of the data. L.K.R-M, J.A.Y, S.D.M., and A.P.L. conceived and designed the study and interpreted the data. L.K.R-M, J.A.Y., J.B., S.Y., K.K.Y.H., M.W., J.P., X.T., M.A., L.B., Z-F.W., and S.L.V. conducted the research. L.K.R-M, J.A.Y., and A.P.L. wrote the manuscript. J.S. and SDM provided critical revisions of the manuscript.

---

**Supplemental Figure 1.**
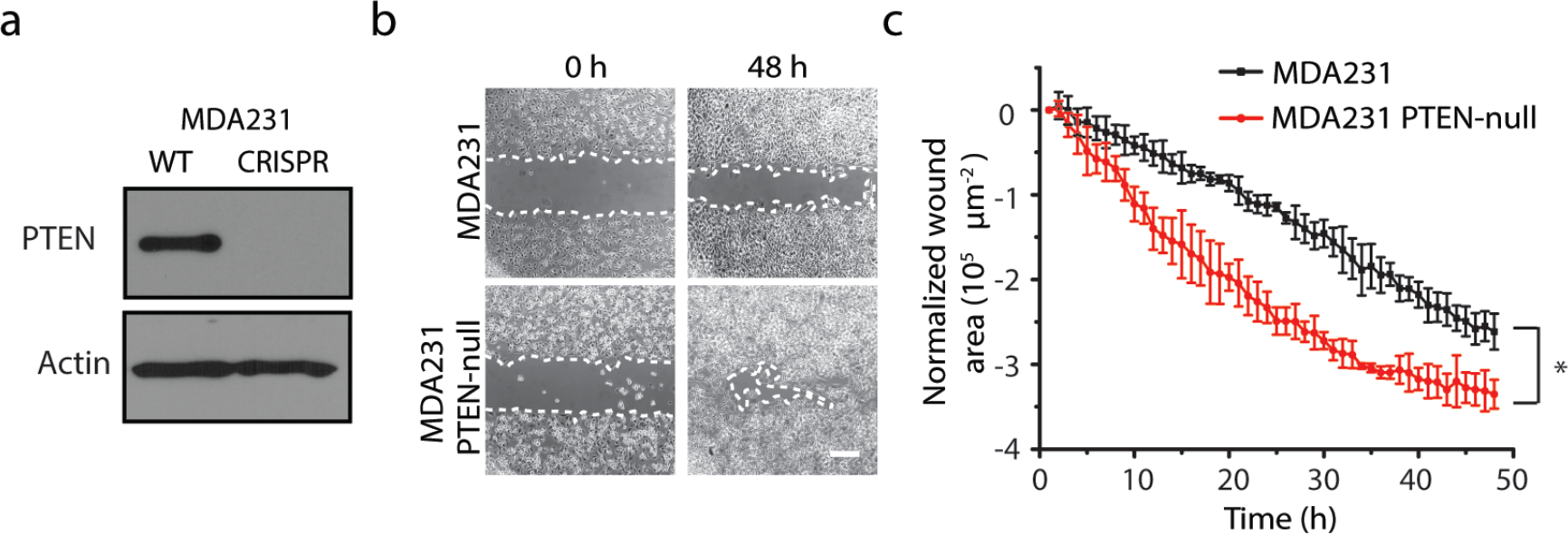
PTEN-null MDA231 cells have lower wound healing rate. **a)** Western blot showing complete knockout of PTEN expression in CRISPR PTEN-null MDA231 cells. **b)** Phase contrast images of a wound healing assay at 0 and 48 hours for MDA231 and MDA231 PTEN-null cells. **c)** Quantification of the wound area as a function of time. The experiment was performed in quadruplicate (mean ± S.D.). * *p* < 0.01 by *t*-test.

**Supplemental Figure 2.**
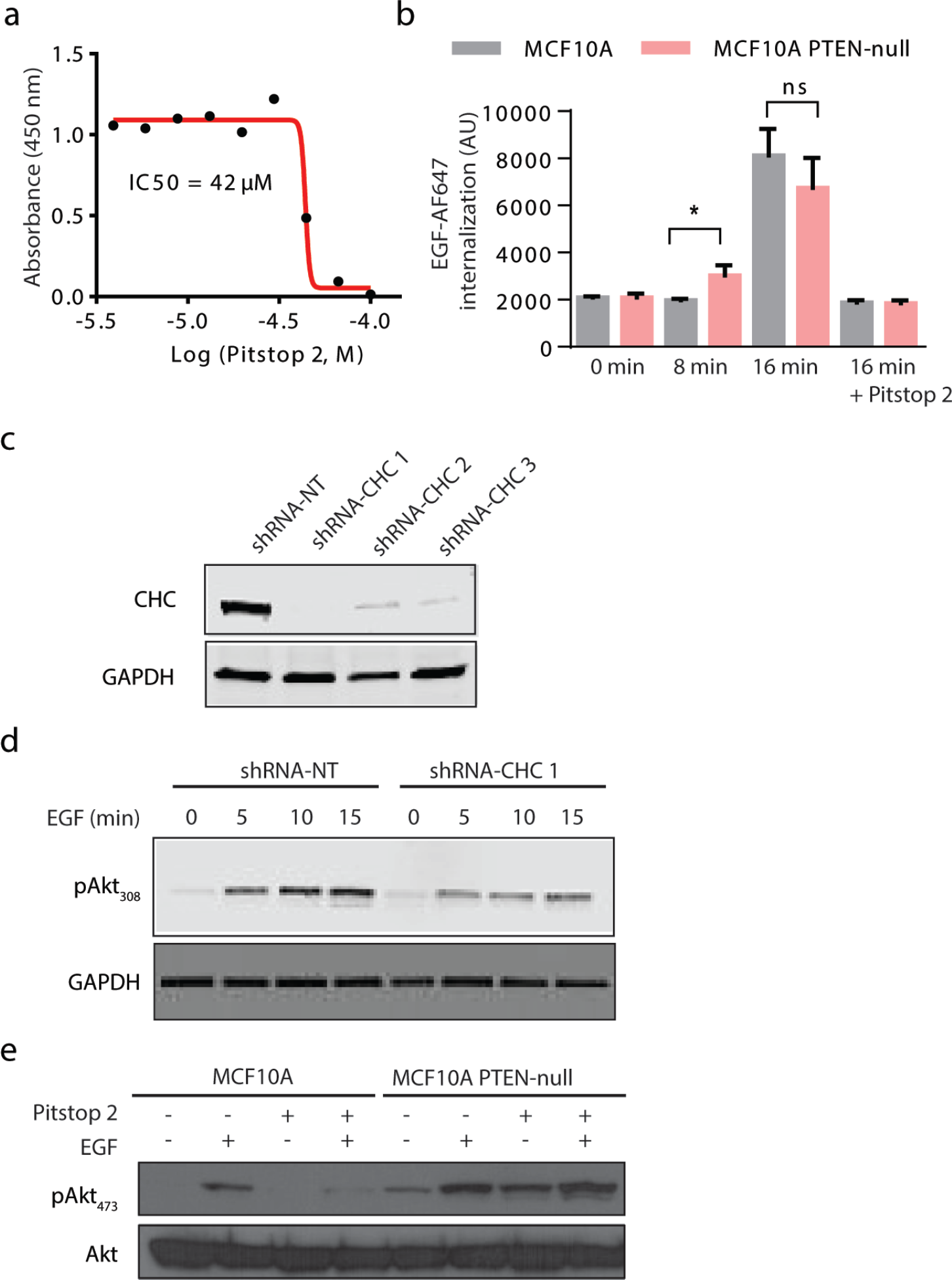
**a)** Determination of the Pitstop 2 cytotoxicity in MCF10A cells by WST-1 assay.The calculated IC50 for Pitstop 2 is 42 µM. **b)** EGF-647 internalization for MCF10A and MCF10A PTEN-null cells stimulated by EGF-647 at different time points with and without Pitstop 2. **c)** MCF10A cells were transduced with lentivirus shRNA targeting clathrin heavy chain (CHC) (sequences 1, 2 and 3). Non-target (NT) sequence was used as a control. **d)** Following transduction, cells were stimulated with 20 ng/mL EGF for indicated time points. Whole-cell lysates were probed with the indicated antibodies. **e)** Western blot analysis of MCF10A and MCF10A PTEN-null cells following EGF stimulation in the presence or absence of Pitstop 2. Whole-cell lysates were probed with the indicated antibodies.

**Supplemental Figure 3.**
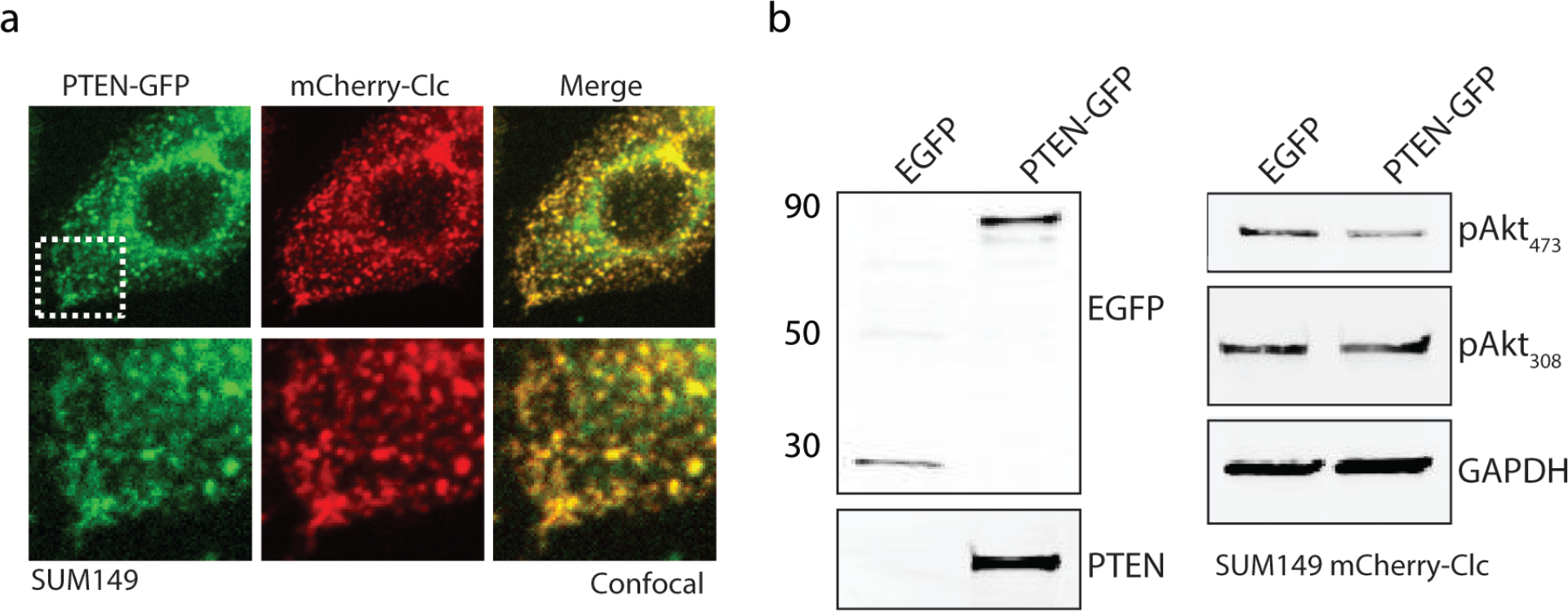
**a)** PTEN-GFP expression extensively co-localizes with mCherry-Clc labeled vesicles in SUM149 mCherry-Clc cells by confocal microscopy. Bottom panels show zoomed-in area marked by the white-dashed square. **b)** Western blot analysis of SUM149 mCherry-Clc cells expressing GFP or PTEN-GFP. Whole-cell lysates were immunobloted with the indicated antibodies.

**Supplemental Figure 4.**
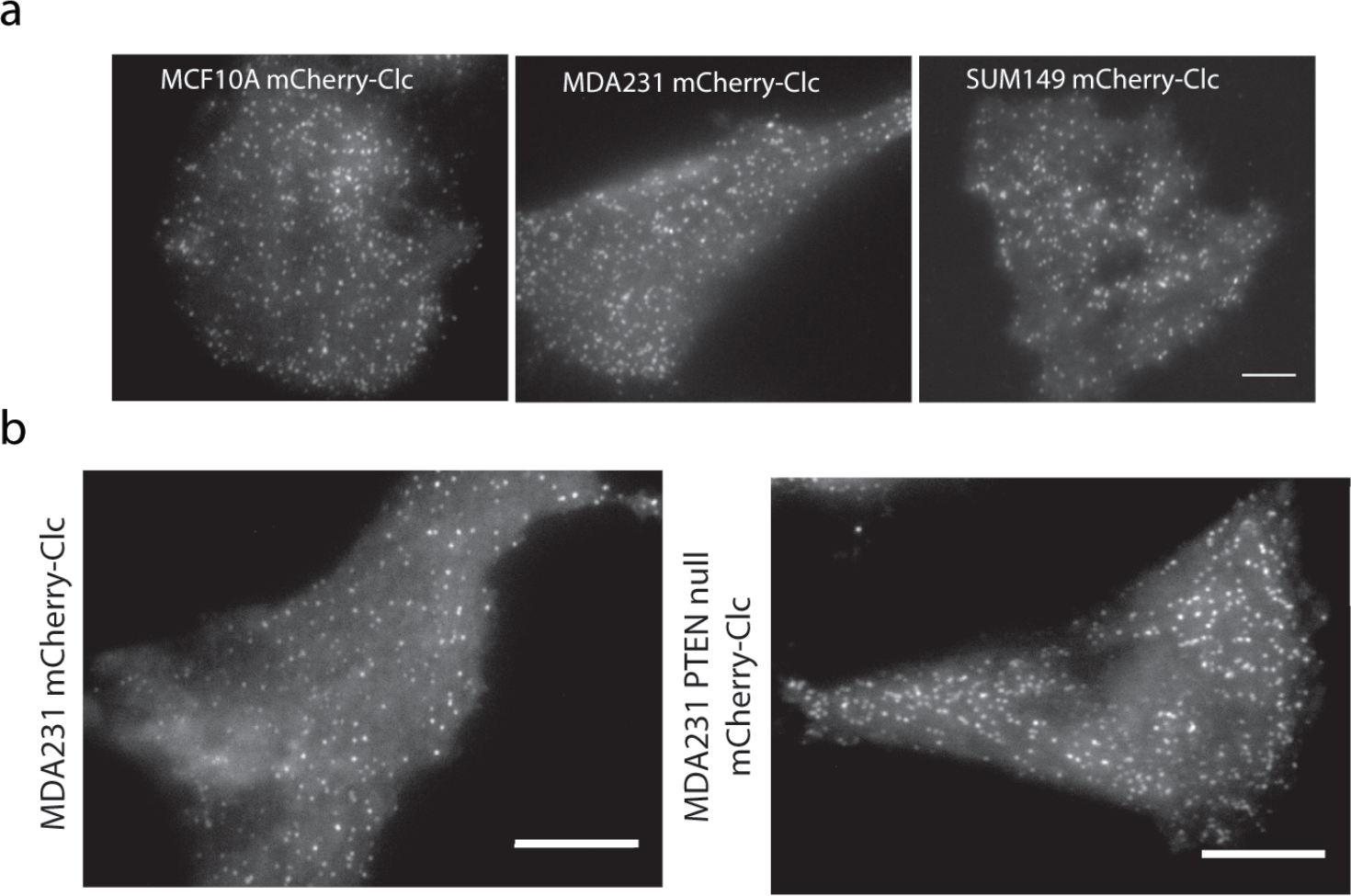
**a)** Representative TIRF microscopy images of cells stably expressing mCherry-Clc in MCF10A, MDA231, and SUM149 cells as indicated. Scale bars is 20 μm. **b)** Representative TIRF images of CCPs for MDA231 stably expressing mCherry-Clc in comparison to CRISPR PTEN-null MDA231 cells. Scale bar is 10 μm.

